## Supplemental Details of the sensitivity analysis for "Sensitivity analysis enlightens effects of connectivity in a Neural Mass Model under Control-Target mode"

### Supplemental Information for the paper: Sensitivity analysis enlightens inhibitory connectivity in a Neural Mass Model under Control-Target mode

Vallet Anaïs[1], Blanco Stéphane[2], Chevallier Coline[2,3], Eustache Francis[1], Gautrais Jacques[2,3], Grandpeix Jean-Yves[4], Joly Jean-Louis[2], Segobin Shailendra[1], Gagnepain Pierre[1],

**1** Normandie Univ, UNICAEN, PSL Research University, EPHE, INSERM, U1077, CHU de Caen, GIP Cyceron, Neuropsychologie et Imagerie de la Mémoire Humaine, 14000 Caen, France

**2** LAPLACE, Université de Toulouse, CNRS, INPT, UPS, Toulouse, France

**3** Centre de Recherches sur la Cognition Animale (CRCA), Centre de Biologie Intégrative (CBI), Université de Toulouse, CNRS, UPS, France

**4** LMD/IPSL, Sorbonne Université, CNRS, École Polytechnique, ENS, Paris, France

*

#### Abstract

This document exposes the formal developments to build sensitivities in the coupled system as a hierarchy of nested sensitivities.

### 1 Sensitivity analysis for the isolated pools

We consider in this section one isolated excitatory pool and one isolated inhibitory pool. Each pool contains an excitatory or inhibitory recurrent coupling and an excitatory forcing. Here, we are interested in the sensitivity of each excitatory and inhibitory pool activity to excitatory forcing.

#### 1.1 Isolated excitatory pool

At the fixed point, the system reads:

$$\begin{cases} 0 = -\beta^E sn^* + \alpha^E T_{glu}(1 - sn^*)rn^* \equiv fn(sn^*, rn^*) \\ rn^* = \dfrac{a_E xn^* - b_E}{1 - e^{-d_E(a_E xn^* - b_E)}} \equiv hn(xn^*) \\ xn^* = W_+ J_{nmda} sn^* + z_E \equiv wn(sn^*, z_E) \end{cases} \tag{1}$$

With the perturbation of $z_E$, we get the new model:

$$\begin{cases} fn(sn^*, rn^*) = 0 \\ rn^* = hn(xn^*) \\ xn^* = wn(sn^*, z_E + \delta z_E) \end{cases} \tag{2}$$

We search the formal expression for the sensitivity $\frac{\delta sn}{\delta z_E}$.
The linearization for the perturbation around fixed point $sn^*$ yields:

$$\begin{cases} \dfrac{\partial fn}{\partial sn}\delta sn + \dfrac{\partial fn}{\partial rn}\delta rn = 0 \\ \delta rn = \dfrac{\mathrm{d}hn}{\mathrm{d}xn}\delta xn \equiv hn'\delta xn \\ \delta xn = \dfrac{\partial wn}{\partial sn}\delta sn + \dfrac{\partial wn}{\partial z_E}\delta z_E \end{cases} \tag{3}$$

Plugging the last two equations into the first, we get:

$$\left(\frac{\partial fn}{\partial sn} + \frac{\partial fn}{\partial rn}\frac{\mathrm{d}hn}{\mathrm{d}xn}\frac{\partial wn}{\partial sn}\right)\delta sn = -\frac{\partial fn}{\partial rn}\frac{\mathrm{d}hn}{\mathrm{d}xn}\frac{\partial wn}{\partial z_E}\delta z_E \tag{4}$$

Rearranging to read how $\delta sn$ depends upon $\delta z_E$:

$$\left(1 + \frac{\partial fn}{\partial sn}^{-1}\frac{\partial fn}{\partial rn}\frac{\mathrm{d}hn}{\mathrm{d}xn}\frac{\partial wn}{\partial sn}\right)\delta sn = -\frac{\partial fn}{\partial sn}^{-1}\frac{\partial fn}{\partial rn}\frac{\mathrm{d}hn}{\mathrm{d}xn}\frac{\partial wn}{\partial z_E}\delta z_E \tag{5}$$

Then we get:

$$\frac{\delta sn}{\delta z_E} = \frac{-\frac{\partial fn}{\partial sn}^{-1}\frac{\partial fn}{\partial rn}\frac{\mathrm{d}hn}{\mathrm{d}xn}\frac{\partial wn}{\partial z_E}}{1 + \frac{\partial fn}{\partial sn}^{-1}\frac{\partial fn}{\partial rn}\frac{\mathrm{d}hn}{\mathrm{d}xn}\frac{\partial wn}{\partial sn}} \tag{6}$$

$$= -\frac{1}{\frac{\partial fn}{\partial sn}\frac{\partial fn}{\partial rn}^{-1}\frac{\mathrm{d}hn}{\mathrm{d}xn}^{-1}\frac{\partial wn}{\partial z_E}^{-1} + \frac{\partial wn}{\partial sn}\frac{\partial wn}{\partial z_E}^{-1}} \tag{7}$$

Considering that

$$\frac{\partial fn}{\partial sn} = -\beta_E - \alpha_E T_{glu} rn^* \tag{8}$$

$$\frac{\partial fn}{\partial rn} = \alpha_E T_{glu}(1 - sn^*) \tag{9}$$

$$\frac{\partial wn}{\partial sn} = W_+ J_{nmda} \tag{10}$$

$$\frac{\partial wn}{\partial z_E} = 1 \tag{11}$$

We can calculate:

$$\frac{\delta sn}{\delta z_E} = \frac{1}{\frac{\beta_E + \alpha_E T_{glu} rn^*}{\alpha_E T_{glu}(1 - sn^*)\frac{\mathrm{d}hn}{\mathrm{d}xn}} - W_+ J_{nmda}} \tag{12}$$

#### 1.2 Isolated inhibitory pool

At the fixed point, the system reads:

$$\begin{cases} 0 = -\beta^I sg^* + \alpha^I T_{gaba}(1 - sg^*) rg^* \equiv fg(sg^*, rg^*) \\ rg^* = \dfrac{a_I xg^* - b_I}{1 - e^{-d_I(a_I xg^* - b_I)}} \equiv hg(xg^*) \\ xg^* = -J_- sg^* + z_I \equiv wg(sg^*, z_I) \end{cases} \tag{13}$$

With the perturbation of $z_I$, we get the new model:

$$\begin{cases} fg(sg^*, rg^*) = 0 \\ rg^* = hg(xg^*) \\ xg^* = wg(sg^*, z_I + \delta z_I) \end{cases} \tag{14}$$

We search the formal expression for the sensitivity $\frac{\delta sg}{\delta z_I}$.
The linearization for the perturbation around fixed point $sg^*$ yields:

$$\begin{cases} \dfrac{\partial fg}{\partial sg}\delta sg + \dfrac{\partial fg}{\partial rg}\delta rg = 0 \\ \delta rg = \dfrac{\mathrm{d}hg}{\mathrm{d}xg}\delta xg \equiv hg' \delta xg \\ \delta xg = \dfrac{\partial wg}{\partial sg}\delta sg + \dfrac{\partial wg}{\partial z_I}\delta z_I \end{cases} \tag{15}$$

Plugging the last two equations into the first, we get:

$$\left(\frac{\partial fg}{\partial sg} + \frac{\partial fg}{\partial rg}\frac{\mathrm{d}hg}{\mathrm{d}xg}\frac{\partial wg}{\partial sg}\right)\delta sg = -\frac{\partial fg}{\partial rg}\frac{\mathrm{d}hg}{\mathrm{d}xg}\frac{\partial wg}{\partial z_I}\delta z_I \tag{16}$$

Rearranging to read how $\delta sg$ depends upon $\delta z_I$:

$$\left(1 + \frac{\partial fg}{\partial sg}^{-1}\frac{\partial fg}{\partial rg}\frac{\mathrm{d}hg}{\mathrm{d}xg}\frac{\partial wg}{\partial sg}\right)\delta sg = -\frac{\partial fg}{\partial sg}^{-1}\frac{\partial fg}{\partial rg}\frac{\mathrm{d}hg}{\mathrm{d}xg}\frac{\partial wg}{\partial z_I}\delta z_I \tag{17}$$

Then we get:

$$\frac{\delta sg}{\delta z_I} = \frac{-\frac{\partial fg}{\partial sg}^{-1}\frac{\partial fg}{\partial rg}\frac{\mathrm{d}hg}{\mathrm{d}xg}\frac{\partial wg}{\partial z_I}}{1 + \frac{\partial fg}{\partial sg}^{-1}\frac{\partial fg}{\partial rg}\frac{\mathrm{d}hg}{\mathrm{d}xg}\frac{\partial wg}{\partial sg}} \tag{18}$$

$$= -\frac{1}{\frac{\partial fg}{\partial sg}\frac{\partial fg}{\partial rg}^{-1}\frac{\mathrm{d}hg}{\mathrm{d}xg}^{-1}\frac{\partial wg}{\partial z_I}^{-1} + \frac{\partial wg}{\partial sg}\frac{\partial wg}{\partial z_I}^{-1}} \tag{19}$$

Considering that

$$\frac{\partial fg}{\partial sg} = -\beta_I - \alpha_I T_{gaba} rg^* \tag{20}$$

$$\frac{\partial fg}{\partial rg} = \alpha_I T_{gaba}(1 - sg^*) \tag{21}$$

$$\frac{\partial wg}{\partial sg} = -J_- \tag{22}$$

$$\frac{\partial wg}{\partial z_I} = 1 \tag{23}$$

We can calculate:

$$\frac{\delta sg}{\delta z_I} = \frac{1}{\frac{\beta_I + \alpha_I T_{gaba} rg^*}{\alpha_I T_{gaba}(1-sg^*)\frac{\mathrm{d}hg}{\mathrm{d}xg}} + J_-} \tag{24}$$

#### 1.3 Functional expressions of the sensitivities for the isolated excitatory and inhibitory pools

We define the function $\varphi n_{_E}$ and $\varphi g_{_I}$ to explicitly express the dependencies of the sensitivities for an isolated excitatory pool and an isolated inhibitory pool:

$$\begin{cases} \varphi n_{_E}(sn^*, z_E) \equiv \left( \dfrac{\beta_E + \alpha_E T_{glu} hn(wn(sn^*, z_E))}{\alpha_E T_{glu}(1 - sn^*) hn'(wn(sn^*, z_E))} - W_+ J_{nmda} \right)^{-1} \\ \varphi g_{_I}(sg^*, z_I) \equiv \left( \dfrac{\beta_I + \alpha_I T_{gaba} hg(wg(sg^*, z_I))}{\alpha_I T_{gaba}(1 - sg^*) hg'(wg(sg^*, z_I))} + J_- \right)^{-1} \end{cases} \tag{25}$$

where

$$h\square'(x) = \frac{(ae^{d(ax-b)}(e^{d(ax-b)} - adx + bd - 1))}{(e^{d(ad-b)} - 1)^2} \tag{26}$$

with $a$, $b$ and $d$ to be taken accordingly.

### 2 Sensitivity analysis for one isolated area

At the fixed point, the system with the two coupled pools reads:

$$\begin{cases} 0 = -\beta^E sn^* + \alpha^E T_{glu}(1 - sn^*) rn^* \equiv fn(sn^*, rn^*) \\ 0 = -\beta^I sg^* + \alpha^I T_{gaba}(1 - sg^*) rg^* \equiv fg(sg^*, rg^*) \end{cases} \tag{27}$$

with

$$\begin{cases} rn^* = \dfrac{a_E xn^* - b_E}{1 - e^{-d_E(a_E xn^* - b_E)}} \equiv hn(xn^*) \\ rg^* = \dfrac{a_I xg^* - b_I}{1 - e^{-d_I(a_I xg^* - b_I)}} \equiv hg(xg^*) \end{cases} \tag{28}$$

in which $xn*$ and $xg*$ represent the respective total input currents:

$$\begin{cases} xn^* = W_+ J_{nmda} sn^* - J_{gaba} sg^* + x_E \\ xg^* = J_{nmda} sn^* - J_- sg^* + x_I \end{cases} \tag{29}$$

where $x_E$ and $x_I$ represent basal forcings (effective external inputs), that we will respectively perturbate.

#### 2.1 Splitting $x_{inter}$ from $x_{intra}$

In order to express closed loop sensitivities of the pools as a function of their open loop sensitivities, we split explicitly the total input currents between an internal current, representing the effect of intra-pool recurrences:

$$\begin{cases} xn^*_{intra} = W_+ J_{nmda} sn^* \equiv wn_{intra}(sn^*) \\ xg^*_{intra} = -J_- sg^* \equiv wg_{intra}(sg^*) \end{cases} \tag{30}$$

and an external current, representing the total amount of forcing, due to the basic external forcing and the feedback current from the alternate pool:

$$\begin{cases} xn^*_{inter} = -J_{gaba} sg^* + x_E \equiv wn_{inter}(sg^*, x_E) \\ xg^*_{inter} = J_{nmda} sn^* + x_I \equiv wg_{inter}(sn^*, x_I) \end{cases} \tag{31}$$

so we have:

$$\begin{cases} xn^* = xn^*_{intra} + xn^*_{inter} \equiv wn(xn^*_{intra}, xn^*_{inter}) \\ xg^* = xg^*_{intra} + xg^*_{inter} \equiv wg(xg^*_{intra}, xg^*_{inter}) \end{cases} \tag{32}$$

#### 2.2 Intermediate variable elimination

We define

$$\vec{s} = \begin{pmatrix} sn^* \\ sg^* \end{pmatrix}, \quad \vec{r} = \begin{pmatrix} rn^* \\ rg^* \end{pmatrix}, \quad \vec{x} = \begin{pmatrix} xn^* \\ xg^* \end{pmatrix}, \quad \overrightarrow{x_{intra}} = \begin{pmatrix} xn^*_{intra} \\ xg^*_{intra} \end{pmatrix}, \quad \overrightarrow{x_{inter}} = \begin{pmatrix} xn^*_{inter} \\ xg^*_{inter} \end{pmatrix} \tag{33}$$

so that we can summarize the fixed point as:

$$\begin{cases} \vec{f}(\vec{s},\vec{r}) = \vec{0} \\ \vec{r} = \vec{h}(\vec{x}) \\ \vec{x} = \vec{w}(\overrightarrow{x_{intra}},\overrightarrow{x_{inter}}) \\ \overrightarrow{x_{intra}} = \overrightarrow{w_{intra}}(\vec{s}) \\ \overrightarrow{x_{inter}} = \overrightarrow{w_{inter}}(\vec{s},x_E,x_I) \end{cases} \tag{34}$$

Plugging intermediate variables $\vec{r}$, $\vec{x}$, $\overrightarrow{x_{intra}}$ into the first equation, Eq 34 can be rewritten as:

$$\begin{cases} \vec{F}(\vec{s},\overrightarrow{x_{inter}}) = \vec{0} \\ \overrightarrow{x_{inter}} = \overrightarrow{w_{inter}}(\vec{s},x_E,x_I) \end{cases} \tag{35}$$

where

$$\begin{cases} Fn(sn^*,xn^*_{inter}) \equiv -\beta^E sn^* + \alpha^E T_{glu}(1-sn^*)\dfrac{a_E(W_+J_{nmda}sn^*+xn^*_{inter})-b_E}{1-e^{-d_E[a_E(W_+J_{nmda}sn^*+xn^*_{inter})-b_E]}} \\ Fg(sg^*,xg^*_{inter}) = -\beta^I sg^* + \alpha^I T_{gaba}(1-sg^*)\dfrac{a_I(-J_-sg^*+xg^*_{inter})-b_I}{1-e^{-d_I[a_I(-J_-sg^*+xg^*_{inter})-b_I]}} \end{cases} \tag{36}$$

and where $\overrightarrow{x_{inter}}$, given by Eq 31, will be the support of information transfer between the two pools (hence, denoted *transfer variables*).

#### 2.3 Expressing sensitivities w.r.t. forcings

We start from Eq 35 where we consider the dependency to $x_E$.

With the perturbation, we get the new model:

$$\begin{cases} \vec{F}(\vec{s},\overrightarrow{x_{inter}}) = \vec{0} \\ \overrightarrow{x_{inter}} = \overrightarrow{w_{inter}}(\vec{s},x_E+\delta x_E,x_I) \end{cases} \tag{37}$$

We search the formal expression for:

$$\begin{pmatrix} \mathscr{A}_{sn,x_E} \\ \mathscr{A}_{sg,x_E} \end{pmatrix} \equiv \begin{pmatrix} \frac{\delta sn}{\delta x_E} \\ \frac{\delta sg}{\delta x_E} \end{pmatrix} = \frac{\overrightarrow{\delta s}}{\delta x_E} \tag{38}$$

The linearization for the perturbation around fixed points $\vec{s^*} = (sn^*, sg^*)$ and $\overrightarrow{x_{inter}} = (xn^*_{inter}, xg^*_{inter})$ yields:

$$\begin{cases} \overline{\overline{\dfrac{\partial F}{\partial s}}}\overrightarrow{\delta s} + \overline{\overline{\dfrac{\partial F}{\partial x_{inter}}}}\overrightarrow{\delta x_{inter}} = \vec{0} \\ \overrightarrow{\delta x_{inter}} = \overline{\overline{\dfrac{\partial w_{inter}}{\partial s}}}\overrightarrow{\delta s} + \overrightarrow{\dfrac{\partial w_{inter}}{\partial x_E}}\delta x_E \end{cases} \tag{39}$$

Plugging the last expression into the first, we get:

$$\overline{\overline{\frac{\partial F}{\partial s}}}\overrightarrow{\delta s} + \overline{\overline{\frac{\partial F}{\partial x_{inter}}\frac{\partial w_{inter}}{\partial s}}}\overrightarrow{\delta s} + \overline{\overline{\frac{\partial F}{\partial x_{inter}}}}\overrightarrow{\frac{\partial w_{inter}}{\partial x_E}}\delta x_E = \vec{0} \tag{40}$$

Rearranging to read how $\overrightarrow{\delta s}$ depends upon $\delta x_E$:

$$(\overline{\overline{\mathbb{I}}} + \overline{\overline{\frac{\partial F}{\partial s}}}^{-1} \overline{\overline{\frac{\partial F}{\partial x_{inter}} \frac{\partial w_{inter}}{\partial s}}}) \overrightarrow{\delta s} = - \overline{\overline{\frac{\partial F}{\partial s}}}^{-1} \overline{\overline{\frac{\partial F}{\partial x_{inter}}}} \overrightarrow{\frac{\partial w_{inter}}{\partial x_E}} \delta x_E \tag{41}$$

Defining

$$\overline{\overline{S}} = - \overline{\overline{\frac{\partial F}{\partial s}}}^{-1} \overline{\overline{\frac{\partial F}{\partial x_{inter}}}} \tag{42}$$

we get:

$$(\overline{\overline{\mathbb{I}}} - \overline{\overline{S}} \overline{\overline{\frac{\partial w_{inter}}{\partial s}}}) \overrightarrow{\delta s} = \overline{\overline{S}} \overrightarrow{\frac{\partial w_{inter}}{\partial x_E}} \delta x_E \tag{43}$$

so that:

$$\begin{pmatrix} \mathscr{A}_{sn,x_E} \\ \mathscr{A}_{sg,x_E} \end{pmatrix} = \overrightarrow{\frac{\delta s}{\delta x_E}} = (\overline{\overline{\mathbb{I}}} - \overline{\overline{S}} \overline{\overline{\frac{\partial w_{inter}}{\partial s}}})^{-1} \overline{\overline{S}} \overrightarrow{\frac{\partial w_{inter}}{\partial x_E}} \tag{44}$$

Now turning to $x_I$, we can proceed the same way, expressing at fixed point the model with pertubation::

$$\begin{cases} \overrightarrow{F}(\overrightarrow{s}, \overrightarrow{x_{inter}}) = \overrightarrow{0} \\ \overrightarrow{x_{inter}} = \overrightarrow{w_{inter}}(\overrightarrow{s}, x_E, x_I + \delta x_I) \end{cases} \tag{45}$$

and we obtain:

$$\begin{pmatrix} \mathscr{A}_{sn,x_I} \\ \mathscr{A}_{sg,x_I} \end{pmatrix} = \overrightarrow{\frac{\delta s}{\delta x_I}} = (\overline{\overline{\mathbb{I}}} - \overline{\overline{S}} \overline{\overline{\frac{\partial w_{inter}}{\partial s}}})^{-1} \overline{\overline{S}} \overrightarrow{\frac{\partial w_{inter}}{\partial x_I}} \tag{46}$$

Using that:

$$\overrightarrow{\frac{\partial w_{inter}}{\partial x_E}} = \begin{bmatrix} 1 \\ 0 \end{bmatrix} \quad \text{and} \quad \overrightarrow{\frac{\partial w_{inter}}{\partial x_I}} = \begin{bmatrix} 0 \\ 1 \end{bmatrix} \tag{47}$$

we finally get a compact expression for the matrix of sensitivities to excitatory perturbations upon external forcings in either excitatory or inhibitory pool:

$$\begin{pmatrix} \mathscr{A}_{sn,x_E} & \mathscr{A}_{sn,x_I} \\ \mathscr{A}_{sg,x_E} & \mathscr{A}_{sg,x_I} \end{pmatrix} = (\overline{\overline{\mathbb{I}}} - \overline{\overline{S}} \overline{\overline{\frac{\partial w_{inter}}{\partial s}}})^{-1} \overline{\overline{S}} \tag{48}$$

##### 2.3.1 Expressing Open Loop Sensitivities

In the closed loop situation, from Eq 31, we have that:

$$\overline{\overline{\frac{\partial w_{inter}}{\partial s}}} = \begin{pmatrix} 0 & -J_{gaba} \\ J_{nmda} & 0 \end{pmatrix} \tag{49}$$

and in the open loop situation, we set: $\overline{\overline{\frac{\partial w_{inter}}{\partial s}}} = \overline{\overline{0}}$.

Under perturbation upon $x_E$, Eq 43 is

$$(\overline{\overline{\mathbb{I}}} - \underbrace{\overline{\overline{S}} \overline{\overline{\frac{\partial w_{inter}}{\partial s}}}}_{\overline{\overline{G}}}) \overrightarrow{\delta s} = \overline{\overline{S}} \overrightarrow{\frac{\partial w_{inter}}{\partial x_E}} \delta x_E \tag{50}$$

where $\overline{\overline{G}}$ is the matrix of feedback gain due to the closed loop between the two pools. In the open loop situation, $\overline{\overline{G}}$ is then nullified while the r.h.s. remains untouched. In this case,

$$\overrightarrow{\delta s}^{O} = \overline{\overline{S}} \overrightarrow{\frac{\partial w_{inter}}{\partial x_E}} \delta x_E \tag{51}$$

represents the effect of perturbation upon fixed point values when the feedback loop has been opened.

The canonical form can then be written as:

$$\left(\overline{\overline{\mathbb{I}}} - \overline{\overline{G}}\right)\overrightarrow{\delta s} = \overrightarrow{\delta s}^{O} \tag{52}$$

We have:

$$\frac{\overrightarrow{\delta s}^{O}}{\delta x_E} = \overline{\overline{S}} \underbrace{\overrightarrow{\frac{\partial w_{inter}}{\partial x_E}}}_{\begin{bmatrix}1\\0\end{bmatrix}} = \begin{pmatrix} \mathscr{A}^{O}_{sn,x_E} \\ 0 \end{pmatrix} \tag{53}$$

Obviously, in the open loop condition, the inhibitory pool is not affected by a perturbation upon $x_E$.

The same way, considering perturbation upon $x_I$, we get:

$$\frac{\overrightarrow{\delta s}^{O}}{\delta x_I} = \overline{\overline{S}} \underbrace{\overrightarrow{\frac{\partial w_{inter}}{\partial x_I}}}_{\begin{bmatrix}0\\1\end{bmatrix}} = \begin{pmatrix} 0 \\ \mathscr{A}^{O}_{sg,x_I} \end{pmatrix} \tag{54}$$

In the open loop condition gain, the excitatory pool is not affected by a perturbation upon $x_I$.

Hence $\overline{\overline{S}}$ reads:

$$\overline{\overline{S}} = \begin{pmatrix} \mathscr{A}^{O}_{sn,x_E} & 0 \\ 0 & \mathscr{A}^{O}_{sg,x_I} \end{pmatrix} \tag{55}$$

and represent the matrix of open loop sensitivities.

###### 2.3.2 Using Isolated Pool Sensitivities

Now, lets consider $z_E$ as:

$$z_E = -J_{gaba} sg^* + x_E = xn_{inter} = wn_{inter}(sg^*) \tag{56}$$

It represents the total amount of forcing at excitatory pool, due to the basic external forcing and the feedback current from the inhibitory pool.

Considering that perturbing here $x_E$ is the same thing as perturbing $z_E$ in the excitatory pool considered as an isolated pool, and that the same is true for the inhibitory pool, we then have

$$\begin{cases} \mathscr{A}^{O}_{sn,x_E} = \varphi n_{_E}(sn^*, -J_{gaba} sg^* + x_E) \\ \mathscr{A}^{O}_{sg,x_I} = \varphi g_{_I}(sg^*, J_{nmda} sn^* + x_I) \end{cases} \tag{57}$$

where $\varphi n_{_E}$ and $\varphi g_{_I}$ are given by Eq 25 and are to be evaluated at the fixed points $sn^*$ and $sg^*$ yielded by the closed loop system, and taking into account the total amount of external forcing.

##### 2.3.3 Closed Loop Sensitivities as functions of Open Loop Sensitivities

We have

$$\overline{\overline{\frac{\partial w_{inter}}{\partial s}}} = \begin{bmatrix} \frac{\partial wn_{inter}}{\partial sn} & \frac{\partial wn_{inter}}{\partial sg} \\ \frac{\partial wg_{inter}}{\partial sn} & \frac{\partial wg_{inter}}{\partial sg} \end{bmatrix} = \begin{bmatrix} 0 & -J_{gaba} \\ J_{nmda} & 0 \end{bmatrix} \tag{58}$$

so, using Eq 55, we get:

$$(\overline{\overline{\mathbb{I}}} - \overline{\overline{S}}\,\overline{\overline{\frac{\partial w_{inter}}{\partial s}}}) = \overline{\overline{\mathbb{I}}} - \begin{bmatrix} \mathscr{A}^O_{sn,x_E} & 0 \\ 0 & \mathscr{A}^O_{sg,x_I} \end{bmatrix} \begin{bmatrix} 0 & -J_{gaba} \\ J_{nmda} & 0 \end{bmatrix} \tag{59}$$

$$= \overline{\overline{\mathbb{I}}} - \begin{bmatrix} 0 & -J_{gaba}\mathscr{A}^O_{sn,x_E} \\ J_{nmda}\mathscr{A}^O_{sg,x_I} & 0 \end{bmatrix} \tag{60}$$

$$= \begin{bmatrix} 1 & J_{gaba}\mathscr{A}^O_{sn,x_E} \\ -J_{nmda}\mathscr{A}^O_{sg,x_I} & 1 \end{bmatrix} \tag{61}$$

Inverting:

$$(\overline{\overline{\mathbb{I}}} - \overline{\overline{S}}\,\overline{\overline{\frac{\partial w_{inter}}{\partial s}}})^{-1} = \frac{1}{1 + J_{gaba}\mathscr{A}^O_{sn,x_E} J_{nmda}\mathscr{A}^O_{sg,x_I}} \begin{bmatrix} 1 & -J_{gaba}\mathscr{A}^O_{sn,x_E} \\ J_{nmda}\mathscr{A}^O_{sg,x_I} & 1 \end{bmatrix} \tag{62}$$

$$\tag{63}$$

Finally,

$$\begin{pmatrix} \mathscr{A}_{sn,x_E} & \mathscr{A}_{sn,x_I} \\ \mathscr{A}_{sg,x_E} & \mathscr{A}_{sg,x_I} \end{pmatrix} = (\overline{\overline{\mathbb{I}}} - \overline{\overline{S}}\,\overline{\overline{\frac{\partial w_{inter}}{\partial s}}})^{-1}\overline{\overline{S}} = \frac{1}{1 + J_{nmda}J_{gaba}\mathscr{A}^O_{sn,x_E}\mathscr{A}^O_{sg,x_I}} \times \begin{bmatrix} \mathscr{A}^O_{sn,x_E} & -J_{gaba}\mathscr{A}^O_{sn,x_E}\mathscr{A}^O_{sg,x_I} \\ J_{nmda}\mathscr{A}^O_{sn,x_E}\mathscr{A}^O_{sg,x_I} & \mathscr{A}^O_{sg,x_I} \end{bmatrix} \tag{64}$$

which expresses closed loop sensitivities (sensitivities for the pools when they are coupled) as functions of open loop sensitivities (sensitivities for the pools with only their recurrent coupling).

#### 2.4 Functional expressions of closed loop sensitivities

We define the functions $\Phi n_{_E}$, $\Phi g_{_E}$, $\Phi n_{_I}$ and $\Phi g_{_I}$ to explicitly express the dependencies of the closed loop sensitivities for an isolated region:

$$
\begin{cases}
\mathscr{A}_{sn,x_E} = \Phi n_{_E}(\overrightarrow{s^*}, x_E, x_I) = \dfrac{\mathscr{A}^O_{sn,x_E}}{1 + J_{nmda} J_{gaba} \mathscr{A}^O_{sn,x_E} \mathscr{A}^O_{sg,x_I}} \\[2ex]
\mathscr{A}_{sg,x_E} = \Phi g_{_E}(\overrightarrow{s^*}, x_E, x_I) = \dfrac{J_{nmda} \mathscr{A}^O_{sn,x_E} \mathscr{A}^O_{sg,x_I}}{1 + J_{nmda} J_{gaba} \mathscr{A}^O_{sn,x_E} \mathscr{A}^O_{sg,x_I}} \\[2ex]
\mathscr{A}_{sn,x_I} = \Phi n_{_I}(\overrightarrow{s^*}, x_E, x_I) = \dfrac{-J_{gaba} \mathscr{A}^O_{sn,x_E} \mathscr{A}^O_{sg,x_I}}{1 + J_{nmda} J_{gaba} \mathscr{A}^O_{sn,x_E} \mathscr{A}^O_{sg,x_I}} \\[2ex]
\mathscr{A}_{sg,x_I} = \Phi g_{_I}(\overrightarrow{s^*}, x_E, x_I) = \dfrac{\mathscr{A}^O_{sg,x_I}}{1 + J_{nmda} J_{gaba} \mathscr{A}^O_{sn,x_E} \mathscr{A}^O_{sg,x_I}}
\end{cases}
\tag{65}
$$

where the open loop sensitivities are given in Eq 57.

### 3 Two coupled area system

We now turn to the coupling between two areas. We seek the expression of their closed loop sensitivities to perturbation as functions of their sensitivities when isolated, as it has been expressed in the previous sections, hence, corresponding to their response to perturbation when the feedback loops are opened at the system scale.

#### 3.1 Splitting $xinter$ from $xintra$

At fixed point, the system can be expressed as:

$$\begin{cases} 0 = -\beta^E sn_i^* + \alpha^E T_{glu}(1 - sn_i^*) rn_i^* & \equiv fn_i(sn_i^*, rn_i^*) \\ 0 = -\beta^I sg_i^* + \alpha^I T_{gaba}(1 - sg_i^*) rg_i^* & \equiv fg_i(sn_i^*, rn_i^*) \end{cases} \tag{66}$$

with

$$\begin{cases} rn_i^* = \dfrac{a_E xn_i^* - b_E}{1 - e^{-d_E(a_E xn_i^* - b_E)}} & \equiv hn_i(xn_i^*) \\ rg_i^* = \dfrac{a_I xg_i^* - b_I}{1 - e^{-d_I(a_I xg_i^* - b_I)}} & \equiv hg_i(xg_i^*) \end{cases} \tag{67}$$

where $i \in \{1, 2\}$.

In Eq 67, $xn_i^*$ and $xg_i^*$ represent the respective total input current to area $i$. In order to express closed loop sensitivities of the areas as a function of their open loop sensitivities, we split explicitly this total input current between an internal current within an area, due to intra-pool recurrence, and coupling between pools:

$$\begin{cases} xn_{intra,i}^* = W_+ J_{nmda} sn_i^* - J_{gaba_i} sg_i^* & \equiv wn_{intra,i}(sn_i^*, sg_i^*) \\ xg_{intra,i}^* = J_{nmda} sn_i^* - J_- sg_i^* & \equiv wg_{intra,i}(sn_i^*, sg_i^*) \end{cases} \tag{68}$$

and external current due to the external inputs and the coupling between the two areas:

$$\begin{cases} xn_{inter,i}^* = k_{E_{ij}} \kappa_{ij} sn_j^* + B_{E_i} & \equiv wn_{inter,i}(sn_j^*) \quad j \neq i \\ xg_{inter,i}^* = (1 - k_{E_{ij}}) \kappa_{ij} sn_j^* + B_{I_i} & \equiv wg_{inter,i}(sn_j^*) \quad j \neq i \end{cases} \tag{69}$$

so that total input currents reads:

$$\begin{cases} xn_i^* = xn_{intra,i}^* + xn_{inter,i}^* = wn_i(xn_{intra,i}^*, xn_{inter,i}^*) \\ xg_i^* = xg_{intra,i}^* + xg_{inter,i}^* = wg_i(xg_{intra,i}^*, xg_{inter,i}^*) \end{cases} \tag{70}$$

#### 3.2 Intermediate variables elimination

We define

$$\vec{s} = \begin{pmatrix} sn_1 \\ sn_2 \\ sg_1 \\ sg_2 \end{pmatrix}, \quad \vec{r} = \begin{pmatrix} rn_1 \\ rn_2 \\ rg_1 \\ rg_2 \end{pmatrix}, \quad \vec{x} = \begin{pmatrix} xn_1 \\ xn_2 \\ xg_1 \\ xg_2 \end{pmatrix}, \quad \overrightarrow{x_{intra}} = \begin{pmatrix} xn_{intra,1} \\ xn_{intra,2} \\ xg_{intra,1} \\ xg_{intra,2} \end{pmatrix}, \quad \overrightarrow{x_{inter}} = \begin{pmatrix} xn_{inter,1} \\ xn_{inter,2} \\ xg_{inter,1} \\ xg_{inter,2} \end{pmatrix} \tag{71}$$

and

$$\overrightarrow{B_E} = \begin{pmatrix} B_{E_1} \\ B_{E_2} \end{pmatrix}, \quad \overrightarrow{B_I} = \begin{pmatrix} B_{I_1} \\ B_{I_2} \end{pmatrix} \tag{72}$$

so that we can summarize the fixed point as:

$$\begin{cases} \overrightarrow{f}(\overrightarrow{s}, \overrightarrow{r}) = \overrightarrow{0} \\ \overrightarrow{r} = \overrightarrow{h}(\overrightarrow{x}) \\ \overrightarrow{x} = \overrightarrow{w}(\overrightarrow{x_{intra}}, \overrightarrow{x_{inter}}) \\ \overrightarrow{x_{intra}} = \overrightarrow{w_{intra}}(\overrightarrow{s}) \\ \overrightarrow{x_{inter}} = \overrightarrow{w_{inter}}(\overrightarrow{s}, \overrightarrow{B_E}, \overrightarrow{B_I}) \end{cases} \tag{73}$$

Plugging intermediate variables $\overrightarrow{r}$, $\overrightarrow{x}$, $\overrightarrow{x_{intra}}$ into the first equation, Eq 73 can be rewritten as:

$$\begin{cases} \overrightarrow{F}(\overrightarrow{s}, \overrightarrow{x_{inter}}) = \overrightarrow{0} \\ \overrightarrow{x_{inter}} = \overrightarrow{w_{inter}}(\overrightarrow{s}, \overrightarrow{B_E}, \overrightarrow{B_I}) \end{cases} \tag{74}$$

where

$$\begin{cases} Fn_i(sn_i^*, sg_i^*, xn_{inter,i}^*) = -\beta^E sn_i^* \\ \qquad + \alpha^E T_{glu}(1 - sn_i^*) \dfrac{a_E(W_+ J_{nmda} sn_i^* - J_{gaba_i} sg_i^* + xn_{inter,i}^*) - b_E}{1 - e^{-d_E[a_E(W_+ J_{nmda} sn_i^* - J_{gaba_i} sg_i^* + xn_{inter,i}^*) - b_E]}} \\ Fg_i(sn_i^*, sg_i^*, xg_{inter,i}^*) = -\beta^I sg_i^* \\ \qquad + \alpha^I T_{gaba}(1 - sg_i^*) \dfrac{a_I(J_{nmda} sn_i^* - J_- sg_i^* + xg_{inter,i}^*) - b_I}{1 - e^{-d_I[a_I(J_{nmda} sn_i^* - J_- sg_i^* + xg_{inter,i}^*) - b_I]}} \end{cases} \tag{75}$$

and where $\overrightarrow{x_{inter}}$, given by Eq 69, will be the transfer variables.

#### 3.3 Expressing sensitivities w.r.t. forcings

##### 3.3.1 Perturbation of one forcing $B$

Here, we build the general expression of the sensitivity, for any given forcing $B$ among $B_{E_1}$, $B_{E_2}$, $B_{I_1}$ or $B_{I_2}$.

The perturbed form of system 74 reads:

$$\begin{cases} \overrightarrow{F}(\overrightarrow{s}, \overrightarrow{x_{inter}}) = \overrightarrow{0} \\ \overrightarrow{x_{inter}} = \overrightarrow{w_{inter}}(\overrightarrow{s}, B + \delta B) \end{cases} \tag{76}$$

By linearization at fixed point, we get:

$$\begin{cases} \overline{\overline{\dfrac{\partial F}{\partial s}}} \overrightarrow{\delta s_B} + \overline{\overline{\dfrac{\partial F}{\partial x_{inter}}}} \overrightarrow{\delta x_{inter\,B}} = \overrightarrow{0} \\ \overrightarrow{\delta x_{inter\,B}} = \overline{\overline{\dfrac{\partial w_{inter}}{\partial s}}} \overrightarrow{\delta s_B} + \overrightarrow{\dfrac{\partial w_{inter}}{\partial B}} \delta B \end{cases} \tag{77}$$

Plugging the second equation into the first, we have:

$$(\overline{\overline{\frac{\partial F}{\partial s}}} + \overline{\overline{\frac{\partial F}{\partial x_{inter}} \frac{\partial w_{inter}}{\partial s}}}) \overrightarrow{\delta s_B} = -\overline{\overline{\frac{\partial F}{\partial x_{inter}}}} \overrightarrow{\frac{\partial w_{inter}}{\partial B}} \delta B \tag{78}$$

Denoting

$$\overline{\overline{S}} = -\overline{\overline{\frac{\partial F}{\partial s}}}^{-1}\overline{\overline{\frac{\partial F}{\partial x_{inter}}}} \tag{79}$$

expression 78 can be written under the canonical form as:

$$(\overline{\overline{\mathbb{I}}} - \overline{\overline{S}}\overline{\overline{\frac{\partial w_{inter}}{\partial s}}})\overrightarrow{\delta s_B} = \overline{\overline{S}}\overrightarrow{\frac{\partial w_{inter}}{\partial B}}\delta B \tag{80}$$

In this expression $\overline{\overline{\frac{\partial w_{inter}}{\partial s}}}$ represents the coupling between the two areas, and is:

$$\overline{\overline{\frac{\partial w_{inter}}{\partial s}}} = \begin{pmatrix} 0 & \frac{\partial wn_{inter,1}}{\partial sn_2} & 0 & 0 \\ \frac{\partial wn_{inter,2}}{\partial sn_1} & 0 & 0 & 0 \\ 0 & \frac{\partial wg_{inter,1}}{\partial sn_2} & 0 & 0 \\ \frac{\partial wg_{inter,2}}{\partial sn_1} & 0 & 0 & 0 \end{pmatrix} = \begin{pmatrix} 0 & k_{E_{12}}\kappa_{12} & 0 & 0 \\ k_{E_{21}}\kappa_{21} & 0 & 0 & 0 \\ 0 & (1-k_{E_{12}})\kappa_{12} & 0 & 0 \\ (1-k_{E_{21}})\kappa_{21} & 0 & 0 & 0 \end{pmatrix} \tag{81}$$

If we set $\overline{\overline{\frac{\partial w_{inter}}{\partial s}}} = \overline{\overline{0}}$ while the r.h.s. remains untouched, we then obtain the effect of perturbation in the open loop case:

$$\overrightarrow{\delta s_B^O} = \overline{\overline{S}}\overrightarrow{\frac{\partial w_{inter}}{\partial B}}\delta B \tag{82}$$

The canonical expression then reads:

$$\boxed{(\overline{\overline{\mathbb{I}}} - \overline{\overline{G}})\overrightarrow{\delta s_B} = \overrightarrow{\delta s_B^O}} \tag{83}$$

where

$$\overline{\overline{G}} = -\overline{\overline{\frac{\partial F}{\partial s}}}^{-1}\overline{\overline{\frac{\partial F}{\partial x_{inter}}\frac{\partial w_{inter}}{\partial s}}} = \overline{\overline{S}}\overline{\overline{\frac{\partial w_{inter}}{\partial s}}} \tag{84}$$

is the feedback gain matrix.

##### 3.3.2 Expressing Open Loop Sensitivities

Perturbations are operated upon transfer variables, given by $\overrightarrow{w_{inter}}$.

Let $\overrightarrow{x_{inter}}^O$ denote the transfer variables in open loop condition when $\overline{\overline{\frac{\partial w_{inter}}{\partial s}}} = \overline{\overline{0}}$ (ie $\kappa_{12} = \kappa_{21} = 0$):

$$\overrightarrow{x_{inter}}^O = \begin{pmatrix} B_{E_1} \\ B_{E_2} \\ B_{I_1} \\ B_{I_2} \end{pmatrix} \tag{85}$$

Since forcing parameters are independent, we have: $\overline{\overline{\frac{\partial w_{inter}}{\partial x^O_{inter}}}} = \overline{\overline{\mathbb{I}}}$.

Hence considering each perturbation one by one, the operator $\overrightarrow{\frac{\partial w_{inter}}{\partial B}}$ acts as a selector of a column of $\overline{\overline{S}}$, so that we can express the sensitivities in closed loop condition as function of sensitivities in open loop condition.

For instance, perturbing $B_{E_1}$, we get:

$$\overrightarrow{\delta s}^O_{B_{E_1}} = \overline{\overline{S}}\,\frac{\overrightarrow{\partial w_{inter}}}{\partial B_{E_1}}\delta B_{E_1} \iff \frac{\overrightarrow{\delta s}^O_{B_{E_1}}}{\delta B_{E_1}} = \overline{\overline{S}}\,\underbrace{\frac{\overrightarrow{\partial w_{inter}}}{\partial B_{E_1}}}_{\begin{bmatrix}1\\0\\0\\0\end{bmatrix}} \tag{86}$$

hence the open loop sensitivities of both areas to a perturbation upon the excitatory pool of the first area yields the first column of $\overline{\overline{S}}$.

Obviously, in the open loop condition, the second area is not perturbed at all.

Furthermore, considering that perturbing here $B_{E_1}$ is the same thing as perturbing $x_{E_1}$ in the first area considered as an isolated area, we have, by definition given in Eq 38, that:

$$\overrightarrow{\delta s}^O_{B_{E_1}} = \begin{pmatrix}\delta sn_1\\ \delta sn_2\\ \delta sg_1\\ \delta sg_2\end{pmatrix}^O_{B_{E_1}} = \begin{pmatrix}\mathscr{A}_{sn_1,x_{E_1}}\delta B_{E_1}\\ 0\\ \mathscr{A}_{sg_1,x_{E_1}}\delta B_{E_1}\\ 0\end{pmatrix} \tag{87}$$

hence the first column of $\overline{\overline{S}}$ is given by:

$$\frac{\overrightarrow{\delta s}^O_{B_{E_1}}}{\delta B_{E_1}} = \begin{pmatrix}\mathscr{A}_{sn_1,x_{E_1}}\\ 0\\ \mathscr{A}_{sg_1,x_{E_1}}\\ 0\end{pmatrix} \tag{88}$$

Following the same lines of reasoning for the three other perturbations, we finally obtain:

$$\overline{\overline{S}} = \begin{pmatrix}\overline{\overline{\mathscr{A}_{sn,x_E}}} & \overline{\overline{\mathscr{A}_{sn,x_I}}}\\ \overline{\overline{\mathscr{A}_{sg,x_E}}} & \overline{\overline{\mathscr{A}_{sg,x_I}}}\end{pmatrix} \tag{89}$$

with

$$\overline{\overline{\mathscr{A}_{sn,x_E}}} = \begin{pmatrix}\mathscr{A}_{sn_1,x_{E_1}} & 0\\ 0 & \mathscr{A}_{sn_2,x_{E_2}}\end{pmatrix}, \qquad \overline{\overline{\mathscr{A}_{sn,x_I}}} = \begin{pmatrix}\mathscr{A}_{sn_1,x_{I_1}} & 0\\ 0 & \mathscr{A}_{sn_2,x_{I_2}}\end{pmatrix} \tag{90}$$

$$\overline{\overline{\mathscr{A}_{sg,x_E}}} = \begin{pmatrix}\mathscr{A}_{sg_1,x_{E_1}} & 0\\ 0 & \mathscr{A}_{sg_2,x_{E_2}}\end{pmatrix}, \qquad \overline{\overline{\mathscr{A}_{sg,x_I}}} = \begin{pmatrix}\mathscr{A}_{sg_1,x_{I_1}} & 0\\ 0 & \mathscr{A}_{sg_2,x_{I_2}}\end{pmatrix} \tag{91}$$

##### 3.3.3 Using Isolated Area Sensitivities

In the same spirit as in Sec 2.3.2, these open loop sensitivities of areas can be expressed by the analytical expression of their sensitivities when considered isolated, yet to be evaluated respectively at fixed points $\overrightarrow{s_1}^*$ and $\overrightarrow{s_2}^*$ yielded by the closed loop system and taking into account the total amount of external forcing, so we write:

$$
\begin{cases}
\mathscr{A}_{sn_i,x_{E_i}} = \Phi n_{_E}(\overrightarrow{s^*} = \overrightarrow{s_i^*}, x_E = xn^*_{inter,i}, x_I = xg^*_{inter,i}) \\
\mathscr{A}_{sg_i,x_{E_i}} = \Phi g_{_E}(\overrightarrow{s^*} = \overrightarrow{s_i^*}, x_E = xn^*_{inter,i}, x_I = xg^*_{inter,i}) \\
\mathscr{A}_{sn_i,x_{I_i}} = \Phi n_{_I}(\overrightarrow{s^*} = \overrightarrow{s_i^*}, x_E = xn^*_{inter,i}, x_I = xg^*_{inter,i}) \\
\mathscr{A}_{sg_i,x_{I_i}} = \Phi g_{_I}(\overrightarrow{s^*} = \overrightarrow{s_i^*}, x_E = xn^*_{inter,i}, x_I = xg^*_{inter,i})
\end{cases}
\tag{92}
$$

where $xn^*_{inter,i}$ and $x_I = xg^*_{inter,i}$ are defined in Eq 69. Analytical expression for the functions $\Phi n_{_E}$, $\Phi g_{_E}$, $\Phi n_{_I}$ and $\Phi g_{_I}$ are explicitly given in section 2.4.

###### 3.3.4 Closed loop sensitivities as functions of Open Loop Sensitivities

From definition 84 for $\overline{\overline{G}}$, we have:

$$
\overline{\overline{G}} = \begin{pmatrix}
\mathscr{A}_{sn_1,x_{E_1}} & 0 & \mathscr{A}_{sn_1,x_{I_1}} & 0 \\
0 & \mathscr{A}_{sn_2,x_{E_2}} & 0 & \mathscr{A}_{sn_2,x_{I_2}} \\
\mathscr{A}_{sg_1,x_{E_1}} & 0 & \mathscr{A}_{sg_1,x_{I_1}} & 0 \\
0 & \mathscr{A}_{sg_2,x_{E_2}} & 0 & \mathscr{A}_{sg_2,x_{I_2}}
\end{pmatrix}
\begin{bmatrix}
0 & k_{E_{12}}\kappa_{12} & 0 & 0 \\
k_{E_{21}}\kappa_{21} & 0 & 0 & 0 \\
0 & (1-k_{E_{12}})\kappa_{12} & 0 & 0 \\
(1-k_{E_{21}})\kappa_{21} & 0 & 0 & 0
\end{bmatrix}
\tag{93}
$$

that we will write as:

$$
\overline{\overline{G}} = \begin{bmatrix}
0 & G_{12} & 0 & 0 \\
G_{21} & 0 & 0 & 0 \\
0 & G_{32} & 0 & 0 \\
G_{41} & 0 & 0 & 0
\end{bmatrix}
\tag{94}
$$

with

$$
\begin{cases}
G_{12} = \mathscr{A}_{sn_1,x_{E_1}} k_{E_{12}}\kappa_{12} + \mathscr{A}_{sn_1,x_{I_1}}(1-k_{E_{12}})\kappa_{12} \\
G_{21} = \mathscr{A}_{sn_2,x_{E_2}} k_{E_{21}}\kappa_{21} + \mathscr{A}_{sn_2,x_{I_2}}(1-k_{E_{21}})\kappa_{21} \\
G_{32} = \mathscr{A}_{sg_1,x_{E_1}} k_{E_{12}}\kappa_{12} + \mathscr{A}_{sg_1,x_{I_1}}(1-k_{E_{12}})\kappa_{12} \\
G_{41} = \mathscr{A}_{sg_2,x_{E_2}} k_{E_{21}}\kappa_{21} + \mathscr{A}_{sg_2,x_{I_2}}(1-k_{E_{21}})\kappa_{21}
\end{cases}
\tag{95}
$$

For the l.h.s. term in expression 83, we then have:

$$
\overline{\overline{\mathbb{I}}} - \overline{\overline{G}} = \left(\begin{array}{cc|cc}
1 & -G_{12} & 0 & 0 \\
-G_{21} & 1 & 0 & 0 \\
\hline
0 & -G_{32} & 1 & 0 \\
-G_{41} & 0 & 0 & 1
\end{array}\right)
\tag{96}
$$

that we will write as:

$$
\overline{\overline{\mathbb{I}}} - \overline{\overline{G}} = \left(\begin{array}{c|c}
\overline{\overline{B_1}} & \overline{\overline{B_2}} \\
\hline
\overline{\overline{B_3}} & \overline{\overline{B_4}}
\end{array}\right)
\tag{97}
$$

Considering the property that

$$\text{If } \overline{\overline{M}} = \begin{pmatrix} \overline{\overline{A}} & \overline{\overline{B}} \\ \overline{\overline{C}} & \overline{\overline{D}} \end{pmatrix} \text{ with } \overline{\overline{D}} \text{ invertible}$$

$$\text{then } \overline{\overline{M}}^{-1} = \begin{pmatrix} \overline{\overline{R}} & \overline{\overline{S}} \\ \overline{\overline{T}} & \overline{\overline{U}} \end{pmatrix} \text{ with } \begin{cases} \overline{\overline{R}} = \left(\overline{\overline{A}} - \overline{\overline{BD}}^{-1}\overline{\overline{C}}\right)^{-1} \\ \overline{\overline{S}} = -\overline{\overline{RBD}}^{-1} \\ \overline{\overline{T}} = -\overline{\overline{D}}^{-1}\overline{\overline{CR}} \\ \overline{\overline{U}} = \overline{\overline{D}}^{-1}\left(\bar{\bar{\mathbb{I}}} - \overline{\overline{CS}}\right) \end{cases}$$

we obtain

$$\left(\bar{\bar{\mathbb{I}}} - \overline{\overline{G}}\right)^{-1} = \left(\begin{array}{c|c} \overline{\overline{C_1}} & \overline{\overline{C_2}} \\ \hline \overline{\overline{C_3}} & \overline{\overline{C_4}} \end{array}\right) \text{ where } \begin{cases} \overline{\overline{C_1}} = \left(\overline{\overline{B_1}} - \overline{\overline{B_2}}(\overline{\overline{B_4}}^{-1})\overline{\overline{B_3}}\right)^{-1} = \overline{\overline{B_1}}^{-1} \\ \overline{\overline{C_2}} = -\overline{\overline{C_1 B_2}}(\overline{\overline{B_4}}^{-1}) = \bar{\bar{0}} \\ \overline{\overline{C_3}} = -\overline{\overline{B_4}}^{-1}\overline{\overline{B_3 C_1}} = -\overline{\overline{B_3 C_1}} = -\overline{\overline{B_3 B_1}}^{-1} \\ \overline{\overline{C_4}} = \overline{\overline{B_4}}^{-1}\left(\bar{\bar{\mathbb{I}}} - \overline{\overline{B_3 C_2}}\right) = \bar{\bar{\mathbb{I}}} \end{cases} \tag{98}$$

$$= \left(\begin{array}{c|c} \overline{\overline{B_1}}^{-1} & \bar{\bar{0}} \\ \hline -\overline{\overline{B_3 B_1}}^{-1} & \bar{\bar{\mathbb{I}}} \end{array}\right) \tag{99}$$

To recover sensitivities for the closed loop condition, we then consider:

$$\left(\bar{\bar{\mathbb{I}}} - \overline{\overline{G}}\right)^{-1}\overline{\overline{S}} = \left(\begin{array}{c|c} \overline{\overline{B_1}}^{-1} & \bar{\bar{0}} \\ \hline -\overline{\overline{B_3 B_1}}^{-1} & \bar{\bar{\mathbb{I}}} \end{array}\right) \left(\begin{array}{c|c} \overline{\overline{\mathscr{A}_{sn,x_E}}} & \overline{\overline{\mathscr{A}_{sn,x_I}}} \\ \hline \overline{\overline{\mathscr{A}_{sg,x_E}}} & \overline{\overline{\mathscr{A}_{sg,x_I}}} \end{array}\right) \tag{100}$$

$$= \left(\begin{array}{c|c} \overline{\overline{B_1}}^{-1}\overline{\overline{\mathscr{A}_{sn,x_E}}} & \overline{\overline{B_1}}^{-1}\overline{\overline{\mathscr{A}_{sn,x_I}}} \\ \hline -\overline{\overline{B_3 B_1}}^{-1}\overline{\overline{\mathscr{A}_{sn,x_E}}} + \overline{\overline{\mathscr{A}_{sg,x_E}}} & -\overline{\overline{B_3 B_1}}^{-1}\overline{\overline{\mathscr{A}_{sn,x_I}}} + \overline{\overline{\mathscr{A}_{sg,x_I}}} \end{array}\right) \tag{101}$$

so that

$$\boxed{\frac{\overrightarrow{\delta s_B}}{\delta B} = \left(\bar{\bar{\mathbb{I}}} - \overline{\overline{G}}\right)^{-1}\overline{\overline{S}}\frac{\overrightarrow{\partial w_{inter}}}{\partial B}} \tag{102}$$

expresses, in full generality for the two-area system, the matrix of sensitivities to a perturbation upon either forcing $B$, and they are expressed as functions of the sensitivities in single-area system, which are in turn expressed as functions of the sensitivities in the single-pool system.

#### 3.4 Sensitivities in a Target-Control system

##### 3.4.1 Extracting the case of interest

We now focus on the question of how sensitivities would drive the response of the excitatory pool of one area to the activation of the excitatory pool of the other one, depending on the connectivity between the two areas. Hence, we attribute role to each area: the area 1 which excitatory pool is positively perturbed will be called ”Control” area (denoted by C), and the area 2 will be called ”Target area” (denoted by T).

From now on, the observable will then be denoted as:

$$\vec{s} = \begin{pmatrix} sn_C \\ sn_T \\ sg_C \\ sg_T \end{pmatrix} \tag{103}$$

and the transfer variables as:

$$\vec{x} = \begin{pmatrix} xn_{intra,C} & = & B_{E_C} & + & k_{E_{CT}}\kappa_{CT} sn_T \\ xn_{intra,T} & = & B_{E_T} & + & k_{E_{TC}}\kappa_{TC} sn_C \\ xg_{intra,C} & = & B_{I_C} & + & (1-k_{E_{CT}})\kappa_{CT} sn_T \\ xg_{intra,T} & = & B_{I_T} & + & (1-k_{E_{TC}})\kappa_{TC} sn_C \end{pmatrix} \tag{104}$$

and we focus on:

$$\begin{pmatrix} \delta sn_T \\ \delta sn_C \end{pmatrix} \tag{105}$$

in response to $\delta B_{E_C}$.

To extract the situation of interest, from the general result above, we then pick the case:

$$\vec{\delta s}_{B_{E_C}} = \left(\overline{\overline{\mathbb{I}}} - \overline{\overline{G}}\right)^{-1} \overline{\overline{S}} \, \overrightarrow{\frac{\partial w_{inter}}{\partial B_{E_C}}} \delta B_{E_C} \tag{106}$$

$$= \left( \begin{array}{c|c} \overline{\overline{B_1}}^{-1} \overline{\overline{\mathscr{A}_{sn,x_E}}} & \overline{\overline{B_1}}^{-1} \overline{\overline{\mathscr{A}_{sn,x_I}}} \\ \hline -\overline{\overline{B_3 B_1}}^{-1} \overline{\overline{\mathscr{A}_{sn,x_E}}} + \overline{\overline{\mathscr{A}_{sg,x_E}}} & -\overline{\overline{B_3 B_1}}^{-1} \overline{\overline{\mathscr{A}_{sn,x_I}}} + \overline{\overline{\mathscr{A}_{sg,x_I}}} \end{array} \right) \begin{pmatrix} 1 \\ 0 \\ \hline 0 \\ 0 \end{pmatrix} \delta B_{E_C} \tag{107}$$

$$= \left( \begin{array}{c} \overline{\overline{B_1}}^{-1} \overline{\overline{\mathscr{A}_{sn,x_E}}} \begin{pmatrix} 1 \\ 0 \end{pmatrix} \\ \hline -\overline{\overline{B_3 B_1}}^{-1} \overline{\overline{\mathscr{A}_{sn,x_E}}} + \overline{\overline{\mathscr{A}_{sg,x_E}}} \begin{pmatrix} 1 \\ 0 \end{pmatrix} \end{array} \right) \delta B_{E_C} \tag{108}$$

$$\equiv \begin{pmatrix} \delta sn_C \\ \delta sn_T \\ \hline \delta sg_C \\ \delta sg_T \end{pmatrix} \tag{109}$$

where the perturbations of interest are in the upper part, and we have:

$$\begin{pmatrix} \delta sn_C \\ \delta sn_T \end{pmatrix} = \overline{\overline{B_1}}^{-1} \overline{\overline{\mathscr{A}_{sn,x_E}}} \begin{pmatrix} 1 \\ 0 \end{pmatrix} \delta B_{E_C} \tag{110}$$

From definition 97, we get

$$\overline{\overline{B_1}}^{-1} = \begin{pmatrix} 1 & -G_{12} \\ -G_{21} & 1 \end{pmatrix}^{-1} = \frac{1}{1 - G_{12}G_{21}} \begin{pmatrix} 1 & G_{12} \\ G_{21} & 1 \end{pmatrix} \tag{111}$$

hence

$$\begin{pmatrix} \delta sn_C \\ \delta sn_T \end{pmatrix} = \frac{1}{1 - G_{12}G_{21}} \begin{pmatrix} 1 & G_{12} \\ G_{21} & 1 \end{pmatrix} \begin{pmatrix} \mathscr{A}_{sn_C, x_{E_C}} & 0 \\ 0 & \mathscr{A}_{sn_T, x_{E_T}} \end{pmatrix} \begin{pmatrix} 1 \\ 0 \end{pmatrix} \delta B_{E_C} \tag{112}$$

$$= \frac{1}{1 - G_{12}G_{21}} \begin{pmatrix} \mathscr{A}_{sn_C, x_{E_C}} & G_{12}\mathscr{A}_{sn_T, x_{E_T}} \\ G_{21}\mathscr{A}_{sn_C, x_{E_C}} & \mathscr{A}_{sn_T, x_{E_T}} \end{pmatrix} \begin{pmatrix} 1 \\ 0 \end{pmatrix} \delta B_{E_C} \tag{113}$$

$$= \frac{1}{1 - G_{12}G_{21}} \begin{pmatrix} \mathscr{A}_{sn_C, x_{E_C}} \\ G_{21}\mathscr{A}_{sn_C, x_{E_C}} \end{pmatrix} \delta B_{E_C} \tag{114}$$

with

$$\begin{cases} G_{12} = \mathscr{A}_{sn_C, x_{E_C}} k_{E_{CT}} \kappa_{CT} + \mathscr{A}_{sn_C, x_{I_C}} (1 - k_{E_{CT}}) \kappa_{CT} \\ G_{21} = \mathscr{A}_{sn_T, x_{E_T}} k_{E_{TC}} \kappa_{TC} + \mathscr{A}_{sn_T, x_{I_T}} (1 - k_{E_{TC}}) \kappa_{TC} \end{cases} \tag{115}$$

For the sake of clarity, $G_{12}$ and $G_{21}$ are denoted $G_{CT}$ and $G_{TC}$ in the main text.
