## Supplemental Set of parameters for "Sensitivity analysis enlightens effects of connectivity in a Neural Mass Model under Control-Target mode"

Table 1: Set of parameters.

| Parameter | Value | Unit |
|---|---|---|
| $\beta^E$ | 6.6 | $\mathrm{s}^{-1}$ |
| $\alpha^E$ | 72 | $s^{-1}mM^{-1}$ |
| $T_{glu}$ | 0.008 | $mM\ s$ |
| $\beta^I$ | 180 | $s^{-1}$ |
| $\alpha^I$ | 530 | $s^{-1}mM^{-1}$ |
| $T_{gaba}$ | 0.003 | $mM\ s$ |
| $a_E$ | 310 | $nC^{-1}$ |
| $b_E$ | 125 | $Hz$ |
| $d_E$ | 0.16 | $s$ |
| $a_I$ | 615 | $nC^{-1}$ |
| $b_I$ | 177 | $Hz$ |
| $d_I$ | 0.087 | $s$ |
| $W_E$ | 1 | — |
| $W_I$ | 0.7 | — |
| $I_0$ | 0.382 | $nA$ |
| $W_+$ | 1.4 | — |
| $J_{nmda}$ | 0.15 | $nA$ |
| $J_{gaba_i}$ | [0.25;3.0] | $nA$ |
| $G$ | 0.69 | — |
| $J_-$ | 1 | $nA$ |
